## Supplementary material for "circHIPK3 nucleates IGF2BP2 and functions as a competing endogenous RNA": Okholm_2024_Supplementary_Figures

Supplementary Figure S1

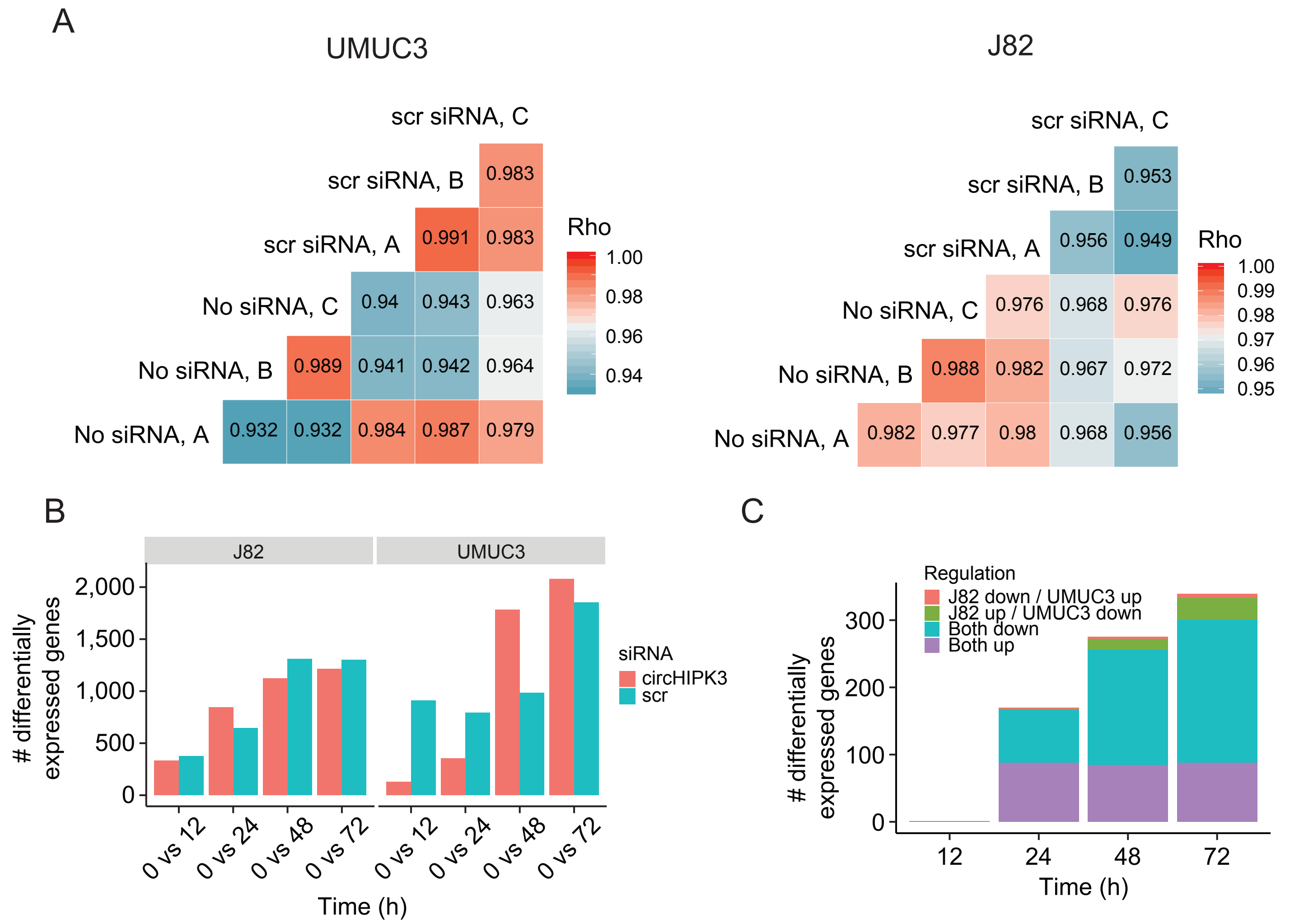


**Supplementary Figure S1: Thousands of genes are deregulated upon circHIPK3 knockdown**
(A) Gene expression correlation upon scr siRNA transfection (scr siRNA, after 1h of incubation) and no transfection (No siRNA) in UMUC3 (left) and J82 (right). Letters (A, B, C) indicate triplicates. Gene expression is highly correlated between scr siRNA samples and No siRNA samples indicating no immediate effect of transfection (Rho > 0.93 for all comparisons, Spearman's rank-order correlation). (B) Number of differentially expressed genes between time points and baseline upon circHIPK3 KD or scr siRNA transfection (Wald test, BH correction with FDR < 0.1). Only genes with perturbed expression profiles across time and/or conditions in UMUC3 (n = 3,072) or J82 (n = 2,389) are considered here. (C) Regulation of shared differentially expressed genes (n = 1,104) between UMUC3 and J82 at each time point. Colours indicate regulation upon circHIPK3 KD vs scr siRNA transfection.

Supplementary Figure S2

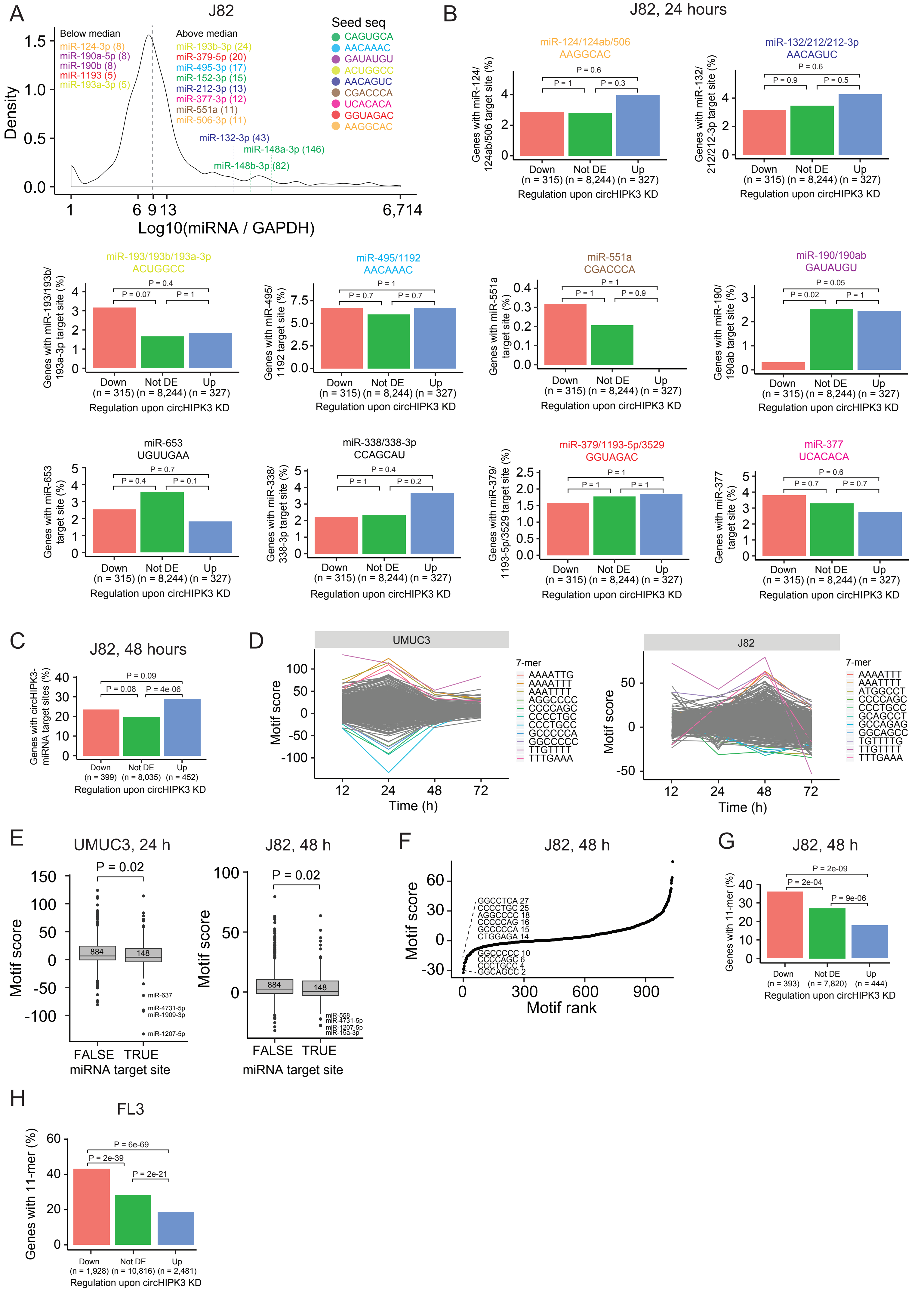

**Supplementary Figure S2: A long motif in circHIPK3 is enriched in downregulated genes upon circHIPK3 knockdown**

(A) miRNA expression profiling in J82 by NanoString. miRNA expression is normalized to GAPDH expression levels. Dotted line indicates median miRNA expression. Conserved miRNAs with target sites in circHIPK3 (circHIPK3-miRNAs) are shown. miRNAs are coloured according to the sequence of their seed site (seed seq). Numbers in parentheses indicate miRNA expression levels. Colors refer to Figure 2A. (B) Percentage of genes in each group with target sites in their 3'UTRs for each circHIPK3-miRNA individually. Gene regulation is based on circHIPK3 KD vs scr siRNA transfection 24 hours post-transfection in J82. Headers indicate the name of the miRNA and the seed site. Colors refer to Figure 2A and Supplementary Figure 2A. The expression of miR-338-3p and miR-653 is not evaluated/detected in J82. P-values obtained by Chi-square Test. (C) Percentage of genes in each group with circHIPK3-miRNA target sites in their 3'UTRs. Gene regulation is based on circHIPK3 KD vs scr siRNA transfection 48 hours post-transfection in J82. P-values obtained by Chi-square Test. (D) Regmex motif scores upon circHIPK3 KD at each time point in UMUC3 (left) and J82 (right). (E) Regmex motif scores for 7-mers corresponding to miRNA target sites (TRUE) and not (FALSE). Numbers correspond to observations in each group. P-values obtained by Wilcoxon Rank Sum Test. (F) Regmex motif scores for circHIPK3-7-mers in J82 (48h). The ten 7-mers with the most negative motif scores in UMUC3 are shown. 7-mers are ranked from most negative to most positive motif scores. Numbers correspond to rank. (G) Percentage of genes in each group containing the 11-mer motif in J82 (48h) and FL3 (24h) cells. P-values obtained by Chi-square Test.

Supplementary Figure S3

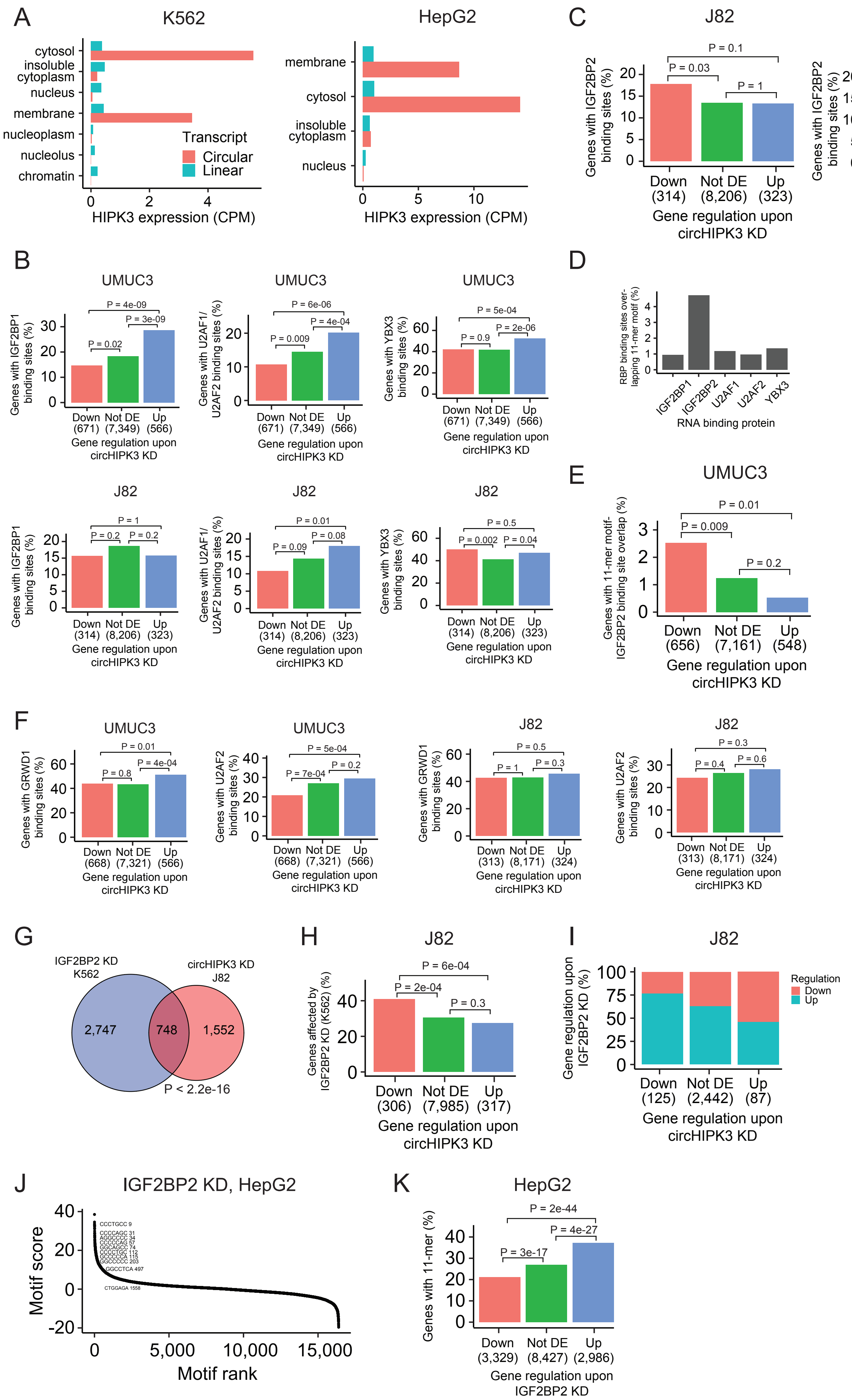

**Supplementary Figure S3: The 11-mer motif in circHIPK3 constitutes a binding site for IGF2BP2**

(A) Expression of circHIPK3 and the corresponding linear transcript in subcellular fractions of K562 (right) and HepG2 (left). CPM = Count per million. (B) Percentage of genes in each group containing RBP binding sites for individual circHIPK3-RBPs in K562. Gene regulation is based on circHIPK3 KD vs scr siRNA transfection 24 hours post-transfection (24h) in UMUC3 (upper panel) and J82 (lower panel). P-values obtained by Chi-square Test. (C) Percentage of genes in each group containing IGF2BP2 binding sites (K562) in J82 (24h) and FL3 cells. P-values obtained by Chi-square Test. (D) Percentage of times circHIPK3-RBP binding sites (K562) overlap the 11-mer motif. (E) Percentage of genes in each group where IGF2BP2 binding sites (K562) overlap the 11-mer motif (UMUC3, 24h). P-values obtained by Chi-square Test. (F) Percentage of genes in each group containing RBP binding sites for individual circHIPK3-RBP in HepG2 (UMUC3 and J82, 24h). P-values obtained by Chi-square Test. (G) Overlap between genes that are affected by circHIPK3 KD in J82 and IGF2BP2 KD in K562. P-value obtained by Fisher's Exact Test. (H) Percentage of genes in each group (J82, 24h) affected by IGF2BP2 KD in K562. P-values obtained by Chi-square Test. (I) Regulation of genes affected by both circHIPK3 KD (J82, 24h) and IGF2BP2 KD (K562). Downregulated genes upon circHIPK3 KD are mainly upregulated upon IGF2BP2 KD and vice versa. (J) Regmex motif enrichment analysis upon IGF2BP2 KD in HepG2. 7-mers are ranked from most positive to most negative motif scores. The ten 7-mers with the most negative motif scores in UMUC3 are shown. Numbers indicate motif rank. All possible 7-mers are evaluated (n = 16,384). (K) Percentage of genes in each group containing the 11-mer motif upon IGF2BP2 KD in HepG2. P-values obtained by Chi-square Test.

#### Supplementary Figure S4

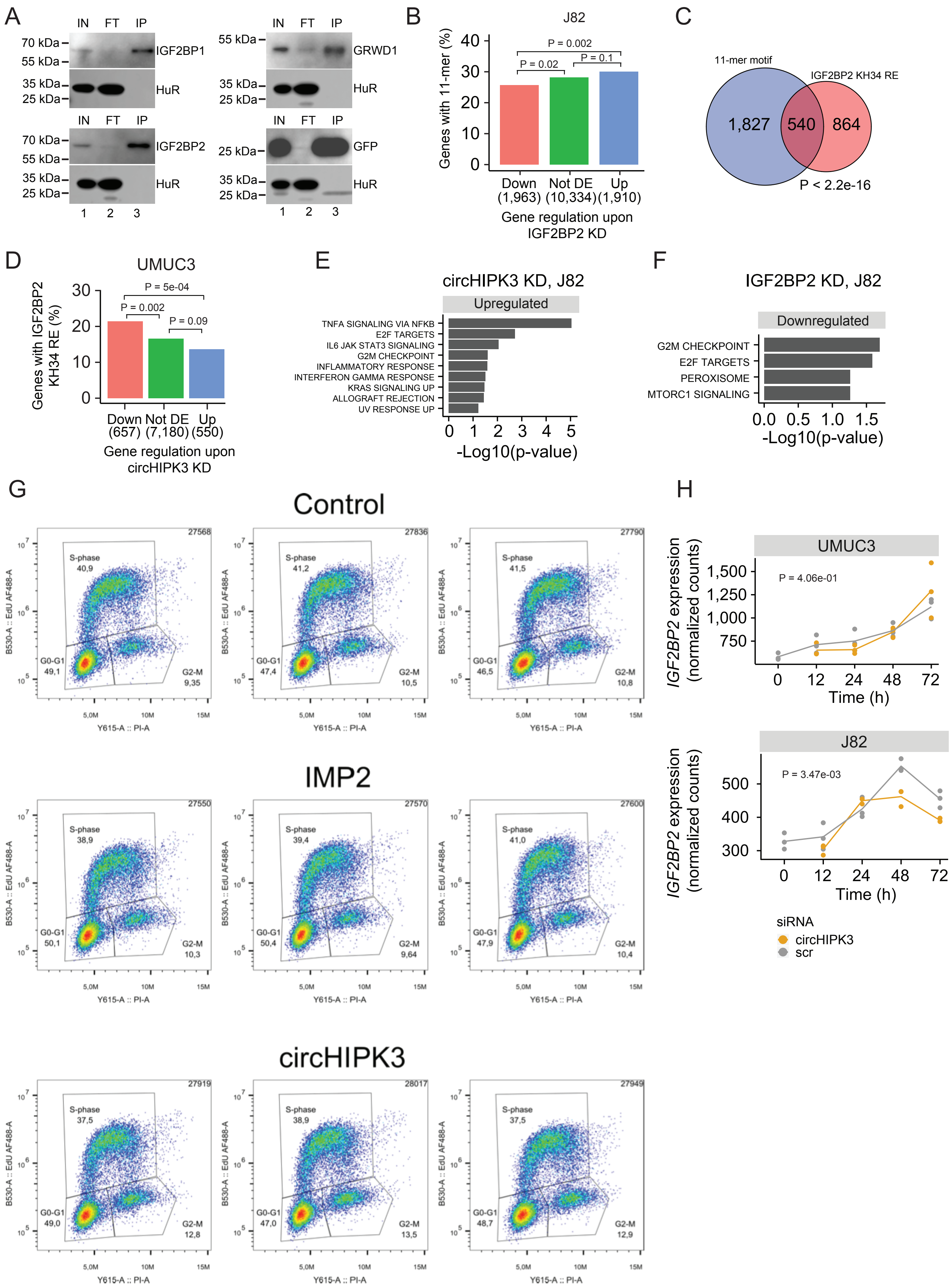

**Supplementary Figure S4: circHIPK3 interacts with IGF2BP2 and affects genes controlling cell cycle progression**
(A) Western blot analyzing protein samples from RNA IP (Figure 4A). Indicated twin-streptagged proteins (”IGF2BP1”, ”IGF2BP2”, ”GRWD1” or ”GFP”) were detected using and anti-streptag antibody. Anti-HuR (”HuR) was used as a loading control. ”IN” = Input, ”FT” = flowthrough, and ”IP” = Immunoprecipitate. (B) Percentage of genes in each group containing the 11-mer motif upon IGF2BP2 KD in J82. P-values obtained by Chi-square Test. (C) Overlap between genes that contain the 11-mer motif and the IGF2BP2 KH34 RE in their 3’UTRs. P-value obtained by Fisher’s Exact Test. (D) Percentage of genes in each group containing the IGF2BP2 KH34 RE upon circHIPK3 KD (UMUC3, 24h). P-values obtained by Chi-square Test. (E) Upregulated hallmarks of cancer upon circHIPK3 KD (48h) in J82 cells (P-value < 0.05 for all shown pathways). (F) Downregulated hallmarks of cancer upon IGF2BP2 KD in J82 (P-value < 0.05 for all shown pathways). (G) Distribution of FL3 cells in the cell cycle. Flow cytometry of cells transiently incubated with EdU followed by AlexaFluor488 labeling (Click-It chemistry). Intensities of propidium iodide (PI) stained DNA (X-axis) and AlexaFluor488 labeled newly synthesized DNA (Y-axis) is plotted. Boxes mark the gates used to estimate the fraction of cells in the indicated phase of the cell cycle. Upper panel: Biological triplicates of control siRNA-treated cells, middle panel: IGF2BP2 (IMP2) knockdown, and lower panel circHIPK3 knockdown. (H) *IGF2BP2* mRNA expression in time-course experiments in UMUC3 (left) and J82 (right). Expression represents DESeq2 normalized counts. P-values obtained by Likelihood Ratio Test.

Supplementary Figure S5

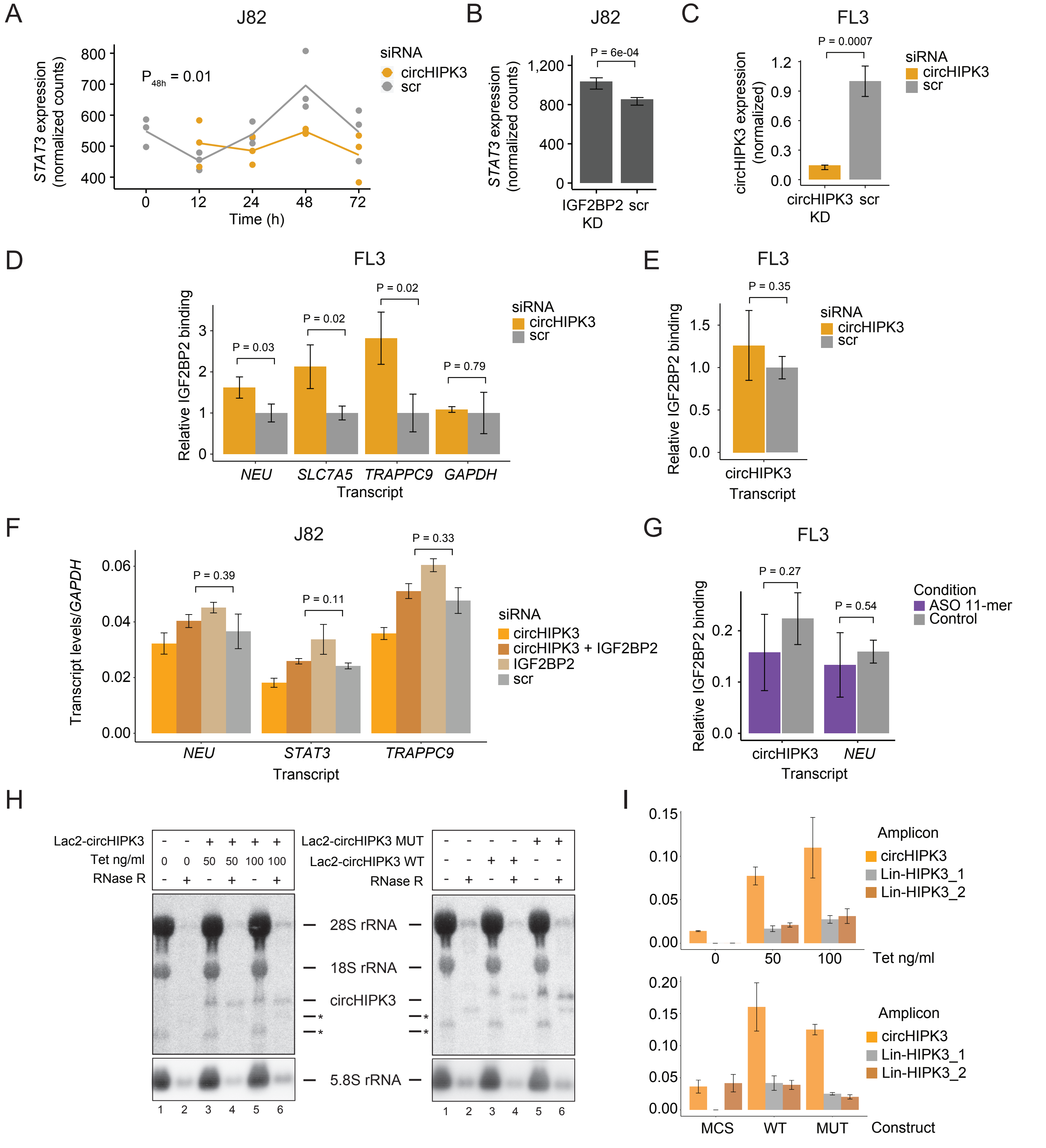


**Supplementary Figure S5: circHIPK3 functions as a competing endogenous RNA for IGF2BP2**
(A) *STAT3* mRNA expression upon circHIPK3 KD in the time-course perturbation experiments in J82 cells. Expression represents DESeq2 normalized counts. P-value (48h) obtained by Wald Test. (B) Expression of *STAT3* upon IGF2BP2 knockdown in J82. P-value obtained by Wald Test. (C) Expression of circHIPK3 upon circHIPK3 KD in FL3 cells. P-values obtained by T-test. (D-E) Relative enrichment of *NEU*, *SLC7A5*, and *TRAPPC9* levels (D) and circHIPK3 (E) between IGF2BP2 IP and input upon circHIPK3 KD. P-values obtained by T-test. Control samples (scr) have been normalized to 1. *GAPDH* was used as a negative control. (F) Rescue experiment showing normalization of target gene expression (*NEU*, *STAT3* and *TRAPPC9*) upon knockdown of both circHIPK3 and IGF2BP2. P-values obtained by T-test. (G) Relative enrichment of circHIPK3 and *NEU* levels between IGF2BP2 IP and input after 11-mer ASO transfection in FL3 cells. (H) Northern blot using RNA from Lac2-circHIPK3 expressing HEK293 FlpIn T-Rex cells or HEK293 cells transfected transiently with Lac2-circHIPK3 expression vectors. * denotes non-specific bands. (I) qRT-PCR using RNA from H circHIPK3 vs linear transcript. Primer efficiencies for amplicons were calculated from standard curves (not shown). (B-G and I) Error bars reflect standard deviation of biological triplicates.

### Supplementary Figure S6

A

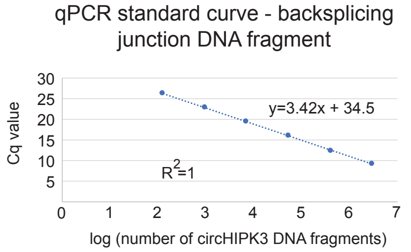


B

5’-CGGCCCATTCTTCAACAAGTGGCCACTGCACTGAAAAAATTGAAAAGTCTTGGTTTAATTCATGCTGA
TCTCAAGCCAGAGAATATTATGTTGGTGGATCCTGTTCGGCAGCCTTACAGGGTTAAAGTAATAGACTTTG
GGTCGGCCAGTCATGTATCAAAGACTGTTTGTTCAACATATCTACAATCTCGGTACTACAG_GTATGGCCT
CACAAGTCTTGGTCTACCCACCATATGTTTATCAAACTCAGTCAAGTGCCTTTTGTAGTGTGAAGAAACTC
AAAGTAGAGCCAAGCAGTTGTGTATTCCAGGAAAGAAACTATCCACGGACCTATGTGAATGGTAGAAACTT
TGGAAATTCTCATCCTCCCACTAAGGGTAGTGCTTTTCAGACAAAGATA-3’

Backsplicing junction fragment

C

| Cells | Cq/10000 | Ave Cq | circHIPK3/PCR | circHIPK3/cell |
|---|---|---|---|---|
| FL3 | 22.9 | 22.74 | 11963.31 | 45 |
| J82 | 22.9 | 22.44 | 13523.67 | 51 |
| UMUC3 | 22.9 | 22.79 | 10593.29 | 40 |

circHIPK3 copies per cell

D

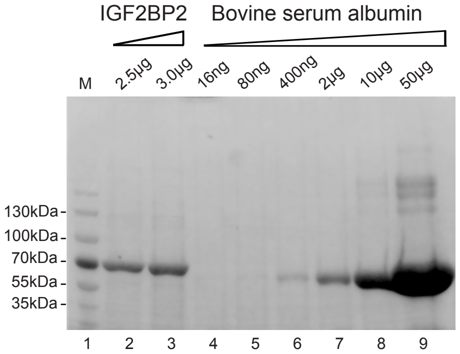


Recombinant IGF2BP2 - Coomassie

E

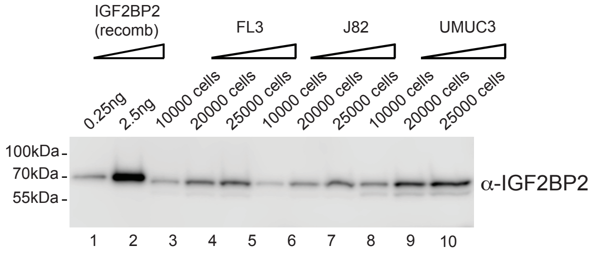


Western blot

F

| Cells | Cells per ng | ng per cell | Moles | IGF2BP2/cell |
|---|---|---|---|---|
| FL3 | 42920 | 2,33E-05 | 3,584E-19 | 215892 |
| J82 | 72686 | 1,38E-05 | 2,117E-19 | 127483 |
| UMUC3 | 35515 | 2,82E-05 | 4,332E-19 | 260903 |

IGF2BP2 copies per cell

**Supplementary Figure S6: Absolute quantification of IGF2BP2 and circHIPK3 in bladder cancer cells.** (A) qPCR standard curve using a DNA fragment spanning the backsplicing junction of circHIPK3. (B) Sequence of DNA fragment used in the qPCR titration experiment. (C) Table summarizing key numbers used to estimate circHIPK3 copy numbers in FL3, J82 and UMUC3 cells (assuming an RT efficiency of 25% compared to direct DNA amplification). Cq/10000 = Cq value for 10.000 molecules. Ave Cq = average Cq value for 3 independent experiments (D) Twin-streptagged-IGF2BP2 was expressed and purified from HEK293-Flp-In T-Rex cells and used for SDS page analysis (lanes 2-3). IGF2BP2 protein levels were quantified by comparison to known amounts of Bovine serum albumin (BSA) using Gelcode Blue. (E) Known amounts of recombinant Twin-streptagged IGF2BP2 were subsequently used for western blotting to estimate levels in FL3, J82 and UMUC3 cells. (F) Table summarizing numbers used to estimate IGF2BP2 abundance in FL3, J82 and UMUC3 cells. *IGF2BP2/Cell* = number of copies of IGF2BP2 per cell in the indicated cell type.
